# Behavioral phenotyping of Zip8 393T-KI mice for *in vivo* study of schizophrenia pathogenesis

**DOI:** 10.1101/2021.10.18.464839

**Authors:** Chantelle E. Terrillion, Byung-hak Kang, Joanna M.P. Melia

## Abstract

Genetic studies have informed on the genetic landscape of schizophrenia, and the next challenge is to link the genetic associations to mechanistic studies. A common single nucleotide polymorphism in the zinc and manganese transporter ZIP8 (rs13107325; ZIP8 A391T) is a top candidate to prioritize for functional studies because it is a missense mutation that results in hypomorphic protein function. With this goal, we have established a mouse model (Zip8 393T-knock-in (KI)), and here, we report the results of brain necropsy and initial behavioral phenotyping experiments in the KI mice using open field testing, elevated plus maze, Y-maze, and trace fear conditioning. Overall, male, homozygous KI mice may exhibit subtle defects in cognition and spatial learning, otherwise the baseline testing supports minimal behavioral differences between wild-type and Zip8 393T-KI mice. There were no genotype-specific alterations of gross or microscopic neuroanatomy. These experiments are important to establish the baseline characteristics of the Zip8 393T-KI mice that may be perturbed in animal models of schizophrenia and position the Zip8 393T-KI mouse as an important model for translational studies of schizophrenia pathogenesis.

## Manuscript

Schizophrenia is a complex, polygenic disease with over 270 genetic loci associated with an increased risk of disease^1^. The genetic associations are dominated by loci in non-coding regions, but, notably, there is a missense polymorphism in *SLC39A8* (rs13107235, ZIP8 A391T), the gene encoding the protein ZIP8. ZIP8 A391T is predicted to be in the top 1.4% of deleterious mutations in the human genome^2^, and it is also one of the most pleiotropic polymorphisms in the human genome^3^. In addition to schizophrenia, ZIP8 A391T is associated with an increased risk of Crohn’s disease^4^, obesity^5^, lower intelligence scores^6^, and alterations in grey matter volume in multiple brain regions^7^, as examples. ZIP8 A391T has a minor allele frequency of approximately 8%^8^. Multiple studies have implicated this polymorphism as the causal variant to account for the association with schizophrenia^1,9^. For these reasons, as we transition from genome wide association studies to functional genomics studies, ZIP8 A391T is a model disease-associated polymorphism to prioritize for functional characterization.

ZIP8 is a metal transporter that participates in manganese homeostasis and zinc transport in the setting of inflammation. ZIP8 is highly conserved across species^10^ with 85% sequence conservation across the entire mouse and human gene with 100% conservation around site 391 (murine equivalent is site 393). We^11^ and others^12^ have taken advantage of this interspecies sequence homology and CRISPR-*Cas9* genome editing to generate a knock-in (KI) mouse model of ZIP8 A391T (murine Zip8 A393T). The Zip8 393T-KI mouse phenocopies ZIP8 391T-associated human phenotypes, including reduced blood Mn, enhanced susceptibility to intestinal inflammation, and altered N-glycosylation^11–13^. The genotypic effect is enhanced in male KI mice, supported by observations in the human population^11,14^. Work-to-date with specific relevance to schizophrenia has suggested ZIP8 A391T may alter glutamate signaling, the neuroinflammatory response, and brain N-glycosylation^13,15^.

To further develop the potential of the Zip8 393T-KI mouse to serve an *in vivo* model to study the link between genetic variation in ZIP8 and schizophrenia risk, we performed baseline behavioral phenotyping and brain necropsy of the KI mice.

Littermate mice from heterozygous Zip8 393T-KI breeding pairs were used for these experiments to yield three genotype groups – wild-type (WT), heterozygous KI, and homozygous KI - of both sexes. Mice were subjected to a series of behavioral phenotyping tests^16,17^ over a period of 8 weeks (**Figure 1; Supplemental Figure 1**). First, to establish equivalency in locomotor activity, open field testing was used. Open field testing can also serve as a screen for anxiety-like behavior where mice prefer to remain in the periphery next to the side wall. In open field testing, total activity and time spent in the center or periphery of the field were equivalent across the three genotype groups in male and female mice. As an additional screen for anxiety-like behavior, the elevated plus maze was performed. No differences were observed between the three genotype groups of either sex. Next, as a test of cognition and spatial learning, the Y-maze test was performed. Here, we observed a possible genotype-driven effect as male, homozygous KI mice spent less time in the novel arm compared to WT male mice, and there was a similar trend in heterozygous mice. These findings met statistical significance by t-test (p=0.0371), but did not meet significance by one-way ANOVA (p=0.1372). In follow-up testing of learning and memory by trace fear conditioning, freezing in response to the context and cue was not changed across the genotype groups. It is important to note that Y-maze and fear conditioning test different aspects of cognition, with Y-maze testing spatial memory while fear conditioning tests associative learning. Therefore, we posit that further testing of more complex spatial learning, like the Morris water maze, may further elucidate defects in cognitive and spatial learning in Zip8 393T-KI male mice.

**Figure 1.**
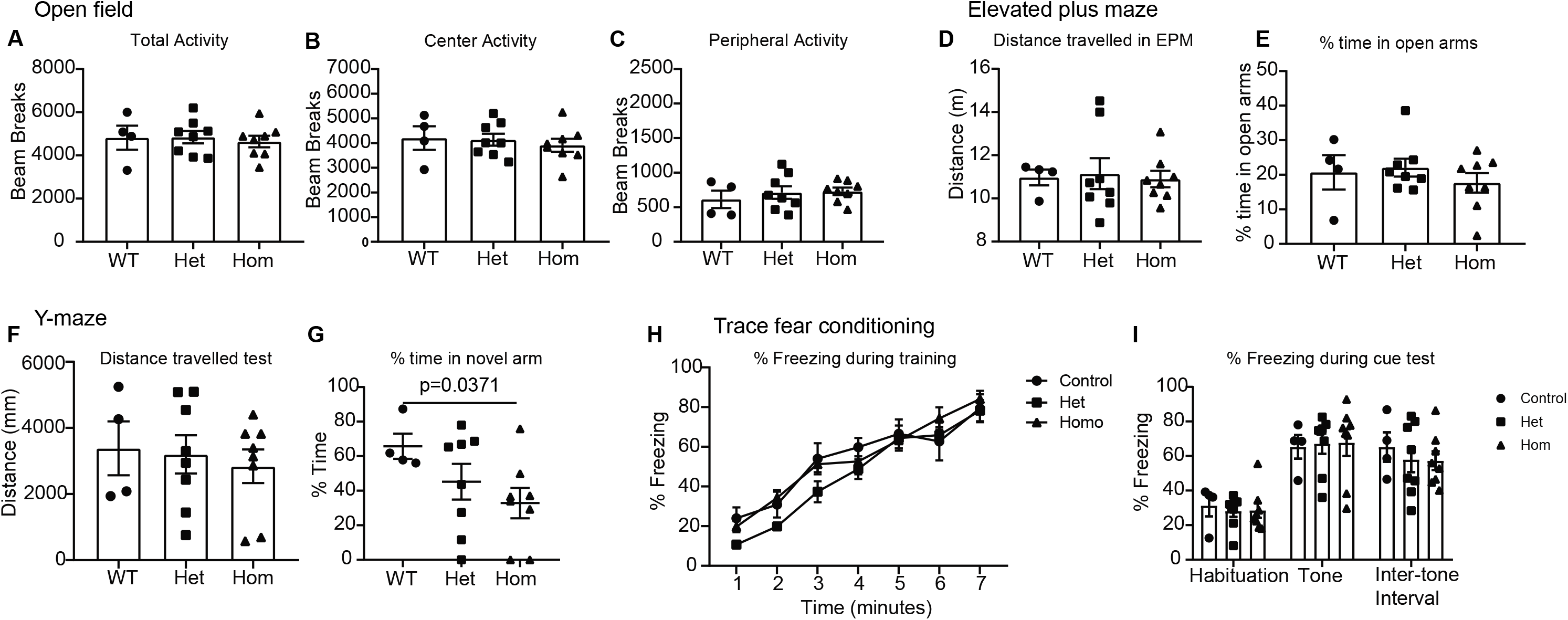
Zip8 393T-KI homozygous male mice exhibit possible defect in cognition and spatial learning, but otherwise normal baseline behavior in motor testing and no phenotype in testing of anxiety behavior and memory. (**A-C**) There are no genotype-specific differences in baseline open-field testing of motor activity with mice of all genotypes showing equal activity levels and time spent in center and periphery. (**D, E**) There are no genotype-specific differences in the elevated plus maze (EPM) testing of anxiety behavior. (**F, G**) Two-step Y-maze tests cognitive and spatial learning. Zip8 393T-KI homozygous mice spent less time in the novel arm (unpaired t-test, p=0.0371); there was a similar trend in the heterozygous mice. One-way ANOVA was not significant (p=0.1372). (**H, I**) There are no genotype-specific differences in trace fear conditioning. Mice were littermates from heterozygous-heterozygous breeding, co-housed, 14-24 weeks of age, n=4-8 male mice/genotype. Mean with S.E.M. represented.

At the end of behavioral testing, formal necropsy was performed for two WT and two homozygous KI mice of each sex with gross and histologic examinations of the brain. The cerebrum, corpus callosum, hippocampus, cerebellum, medulla, ventricles, and pituitary were all determined to be within normal limits upon review by a veterinary pathologist. We suspect genotype-driven differences in grey matter volume, as associated with ZIP8 391-Thr in the UK Biobank study ^7^, will require magnetic resonance imaging studies in the KI mice.

Taken together, these initial experiments are important to establish baseline behavioral characteristics and neuroanatomy in the Zip8 393T-KI mice. The overall, near equivalency across genotypes in these initial experiments make it possible to use these mice in animal models related to schizophrenia, like drug-induced models^18^, and support further testing of more complex spatial learning. We position the Zip8 393T-KI mouse as an important model for translational studies of schizophrenia pathogenesis.

## Methods

### Animal care

Description of generation of the KI mice is as published^11^. Mice were derived from heterozygous breeding pairs. Mice were co-housed by sex and exposed to a 12 hour light-dark cycle. Mice were fed Harlan Teklad Global 18% Protein Extruded Diet 2018SX that includes 100PPM Mn. Care and experimental procedures were approved by the Johns Hopkins University Animal Care and Use Committee.

### Open field

Locomotor activity was assessed over 30 minutes in a 40×40cm activity chamber with infrared beams (San Diego Instruments Inc., San Diego, CA, USA). Horizontal activity, as well as time spent in the center or periphery of the chamber, was automatically recorded.

### Y-Maze spontaneous alternation

Working memory was assessed in the Y-maze spontaneous alternation task. Mice were placed at the end of one arm of a Y-maze consisting of three 38cm long arms (San Diego Instruments Inc., San Diego, CA, USA) and allowed to explore the maze for five minutes. Distance travelled and entries into the arms were automatically recorded using Anymaze Tracking Software (Stoelting, Co., Wood Dale, IL, USA). The percent alternation was calculated using the equation % Alternation = (correct alternations/ (total arm entries – 2))*100.

### Y-Maze spatial recognition

Spatial memory was assessed using the Y-maze spatial recognition test. This test consists of a “Training” phase and a “Test” phase. During the “Training” phase, one arm of a Y-maze consisting of three 38cm long arms (San Diego Instruments) was blocked off. Mice were placed at the end of one of two open arms and allowed to explore for five minutes. After a 30-minute inter-trial interval, the “Test” phase began. The blockade was removed, and mice were allowed to explore all three arms of the maze for five minutes. Distance travelled and time spent in each arm was automatically recorded using Cleversys Topscan tracking software (Cleversys, Inc, Reston, VA, USA). Data from the first two minutes of the “Test” phase were used to evaluate percent time spent in the novel arm.

### Trace fear conditioning

Trace fear conditioning was conducted as previously described in Terrillion et al^16^. Briefly, trace fear conditioning consisted of a habituation day, a training day, and a test day over three consecutive days. On the habituation day, mice were exposed to the shock box (Coulbourn, Holliston, MA, USA) for ten minutes. On the training day, mice were placed in the shock box and given a two-minute habituation, after which a 20-second white noise tone (80db, 2000 Hz) was delivered. Twenty seconds following the termination of the tone, a scrambled two-second 0.5mA shock was delivered. The tone-shock pairing was repeated three additional times. On the test day, mice were placed in the shock box for three minutes to measure freezing in response to the context. Mice were then placed in a separate context and freezing in response to the 20-second white noise tone was measured. Freezing behavior was automatically scored using Cleversys Freezescan (Cleversys Inc., Reston, VA, USA).

### Necropsy

Mice were fasted overnight, individually euthanized by exposure to gradually increasing concentrations of CO2, ana perfused with heparinized saline followed by 10% NBF via left cardiac ventricle. Perfused tissues were fixed in 10% NBF for at least 24 h prior to trimming. The head and hindlimb were separately decalcified with Formical 4 (StatLab Medical Products). Gross examination and tissue collection was performed as described in Brayton, et al^19^. Sections (~5 μm) were stained with hematoxylin and eosin and reviewed by a veterinary pathologist (B. Kang).

### Statistical analysis

Genotype-specific effect was tested using t-test or one-way ANOVA in Prism 8.0 (Graphpad). Sample size and statistical test indicated in figure legend.

## Data availability

Data are available upon request.

## Acknowledgments

This work was supported through NIH (DK114478 to J. M.) and internal funding from the Johns Hopkins University School of Medicine as part of the 2017 Core Coins program. We appreciated the support of the Behavioral Core of the School of Medicine, as well as helpful discussions with past director, Dr. Mikhail Pletnikov, MD, PhD.

## Author Contributions

The authors confirm contribution to the paper as follows: study conception and design: CT, BK, JM; data collection: CT, BK; analysis and interpretation of the results: CT, BK, JM; draft manuscript preparation: CT, JM. All authors reviewed the results and approved the final version of the manuscript.

## Competing Interest Statement

The authors have no competing interests to declare.

## Notes

### Competing Interest Statement

The authors have declared no competing interest.

