## Supplemental figure 1 for "Behavioral phenotyping of Zip8 393T-KI mice for *in vivo* study of schizophrenia pathogenesis"

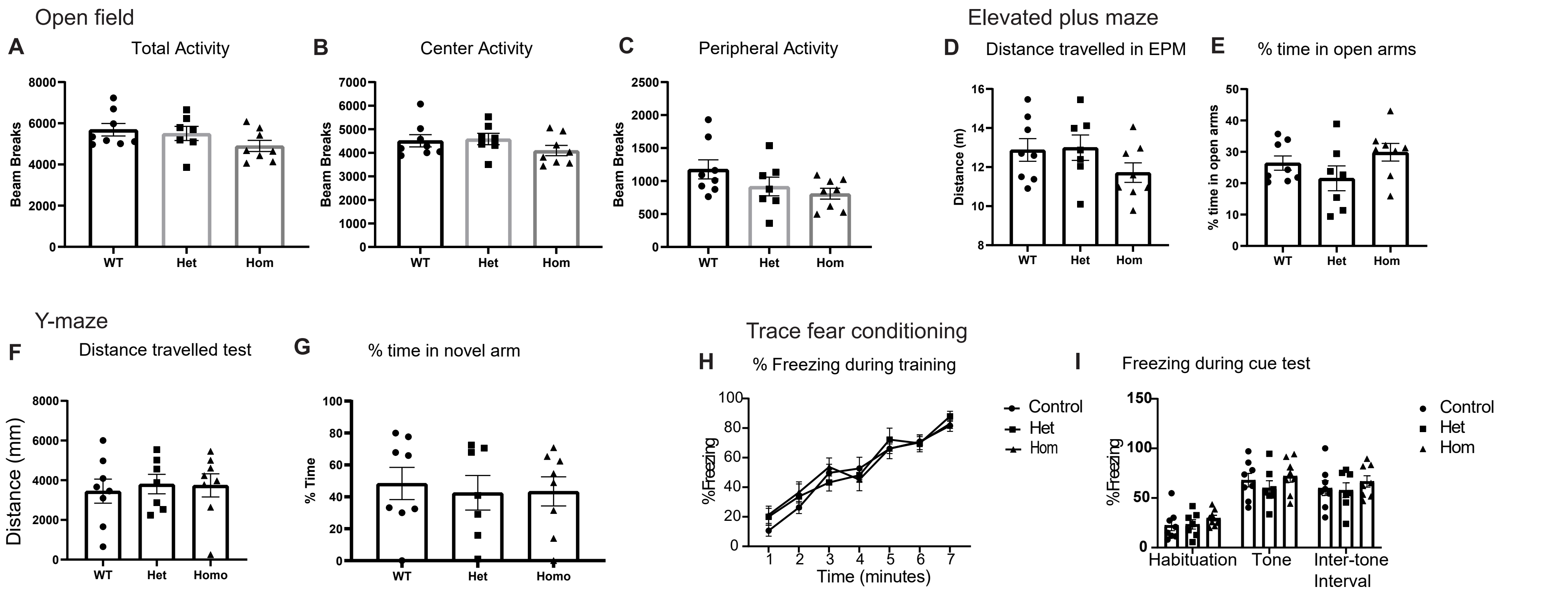


**Supplemental Figure 1. Female Zip8 393T-KI mice exhibit normal baseline behavior in motor testing and no phenotype in testing of anxiety behavior, memory, cognitive and spatial learning.** (**A-C**) There are no genotype-specific differences in baseline open-field testing of motor activity with mice of all genotypes showing equal activity levels and time spent in center and periphery. (**D, E**) There are no genotype-specific differences in the elevated plus maze (EPM) testing of anxiety behavior. (**F, G**) There are no genotype-specific differences in the two-step Y-maze tests of cognitive and spatial learning. (**H, I**) There are no genotype-specific differences in trace fear conditioning. Mice were littermates from heterozygous-heterozygous breeding, co-housed, 14-24 weeks of age, n=4-8 female mice/genotype. Mean with S.E.M. represented.
